# Targeted antifungal liposomes

**DOI:** 10.1101/518043

**Authors:** Suresh Ambati, Aileen R. Ferarro, S. Earl Khang, Xiaorong Lin, Michelle Momany, Zachary Lewis, Richard B. Meagher

**Affiliations:** Department of Genetics, University of Georgia, Athens, GA. 30602; Department of Microbiology, University of Georgia, Athens, GA. 30602; Fungal Biology Group and Department of Plant Biology, University of Georgia, Athens, GA. 30602

## Abstract

*Aspergillus species* cause pulmonary invasive aspergillosis resulting in nearly a hundred thousand deaths each year. Patients at the greatest risk of developing life-threatening aspergillosis have weakened immune systems and/or various lung disorders. Patients are treated with antifungals such as amphotericin B (AmB), casofungin acetate, or triazoles (itraconazole, voriconazole etc.), but these antifungal agents have serious limitations due to lack of sufficient fungicidal effect and human toxicity. Liposomes with AmB intercalated into the lipid membrane (AmBisomes, AmB-LLs), have several-fold reduced toxicity compared to detergent solubilized drug. However, even with the current antifungal therapies, one-year survival among patients is only 25 to 60%. Hence, there is a critical need for improved antifungal therapeutics.

Dectin-1 is a mammalian innate immune receptor in the membrane of some leukocytes that binds as a dimer to beta-glucans found in fungal cell walls, signaling fungal infection. Using a novel protocol, we coated AmB-LLs with Dectin-1’s beta-glucan binding domain to make DEC-AmB-LLs. DEC-AmB-LLs bound rapidly, efficiently, and with great strength *Aspergillus fumigatus* and to *Candida albicans* and *Cryptococcus neoformans*, highly divergent fungal pathogens of global importance. By contrast, un-targeted AmB-LLs and BSA-coated BSA-AmB-LLs showed 200-fold lower affinity for fungal cells. DEC-AmB-LLs reduced the growth and viability of *A. fumigatus* an order of magnitude more efficiently than untargeted control liposomes delivering the same concentrations of AmB, in essence increasing the effective dose of AmB. Future efforts will focus on examining pan-antifungal targeted liposomal drugs in animal models of disease.

**Tweet:** We coated anti-fungal drug loaded liposomes to fungal cell walls with a beta-glucan binding protein and thereby increased drug effectiveness by an order of magnitude.

**Importance:** The fungus *Aspergillus fumigatus* causes pulmonary invasive aspergillosis resulting in nearly a hundred thousand deaths each year. Patients are often treated with antifungal drugs such as amphotericin B loaded into liposomes, AmB-LLs, but all antifungal drugs including AmB-LLs have serious limitations due to human toxicity and insufficient fungal cell killing. Even with the best current therapies, one-year survival among patients with invasive aspergillosis is only 25 to 60%. Hence, there is a critical need for improved antifungal therapeutics.

Dectin-1 is a mammalian protein that binds to beta-glucan polysaccharides found in nearly all fungal cell walls. We coated AmB-LLs with Dectin-1 to make DEC-AmB-LLs. DEC-AmB-LLs bond strongly to fungal cells, while AmB-LLs had little affinity. DEC-AmB-LLs killed or inhibited *A. fumigatus* ten times more efficiently than untargeted lipsomes, increasing the effective dose of AmB. Dectin-1 coated liposomes targeting fungal pathogens have the potential to greatly enhance antifungal therapeutics.

## Introduction

### Invasive fungal infections

Hundreds of species of indigenous fungi cause a wide variety of diseases including aspergillosis, blastomycosis, candidiasis, cryptococcosis, coccidioidomycosis (valley fever), and Pneumocystis pneumonia (PCP). Collectively pathogenic fungi infect many different organs, but lungs are the most common site for deep mycoses. Globally aspergillosis, candidiasis, and cryptococcosis kill about one million or more people each year (1, 2).

*Aspergillus fumigatus* and related *Aspergillus species* cause aspergillosis (2). Patients at the greatest risk of developing life-threatening aspergillosis have weakened immune systems, for example, from stem cell transplants or organ transplants or have various lung diseases, including tuberculosis, chronic obstructive pulmonary disease, cystic fibrosis, or asthma. Among immunocompromised patients, aspergillosis is the second most common fungal infection, after candidiasis (3, 4). Additional costs associated with treating invasive aspergillosis are estimated at $40,000 per child and $10,000 per adult. Patients with aspergillosis are treated with antifungals such as amphotericin B, caspofungin or triazoles. Even with antifungal therapy, however, one-year survival among immunocompromised patients with aspergillosis is only 25 to 60%. Furthermore, all known antifungal agents that treat aspergillosis are quite toxic to human cells(5, 6). The goal of our research has been to develop a targeted liposomal strategy that improves antifungal drug delivery and enhances therapeutic efficacy.

### Liposomal AmB

Amphotericin B (AmB) is the most commonly used agent for many kinds of fungal infections, including aspergillosis. Because AmB binds the fungal plasma membrane sterol ergosterol more efficiently than the mammalian sterol cholesterol, AmB is more toxic to fungal cells. The side effects of Amphotericin B include neurotoxicity and/or nephrotoxicity and/or hepatoxicity (5, 6) and can result in death of the patient (1).

Amphotericin B loaded liposomes, AmB-LLs, penetrate more efficiently to various organs (7, 8), penetrate the cell wall (9) and show reduced toxicity at higher, more effective doses of AmB than the second most commonly used AmB product, deoxycholate detergent-solubilized AmB (5, 6, 10, 11). AmB is an amphipathic molecule. Its long lipophilic polyene end intercalates into the lipid bilayer of liposomes, while its hydrophilic end is positioned on the liposomal surface as modeled in **Fig. 1**. Commercial untargeted spherical AmB-LLs are called AmBisomes (12, 13). However, AmB-LLs still produce AmB human toxicity, such as renal toxicity in 50% of patients (5, 6, 11). When infected mice are treated with AmB-LLs, viable numbers of *A. fumigatus* cells in homogenized lung tissue were only reduced by 70% (14, 15), leaving large fungal cell populations behind. This large residual fungal population may result in recurrence and subsequent mortality after treatment. We explored the targeting of AmB-LLs to *Aspergillus fumigatus* cells to meet the pressing need to improve the quality of antifungal drug formulations (1).

**Fig. 1.**
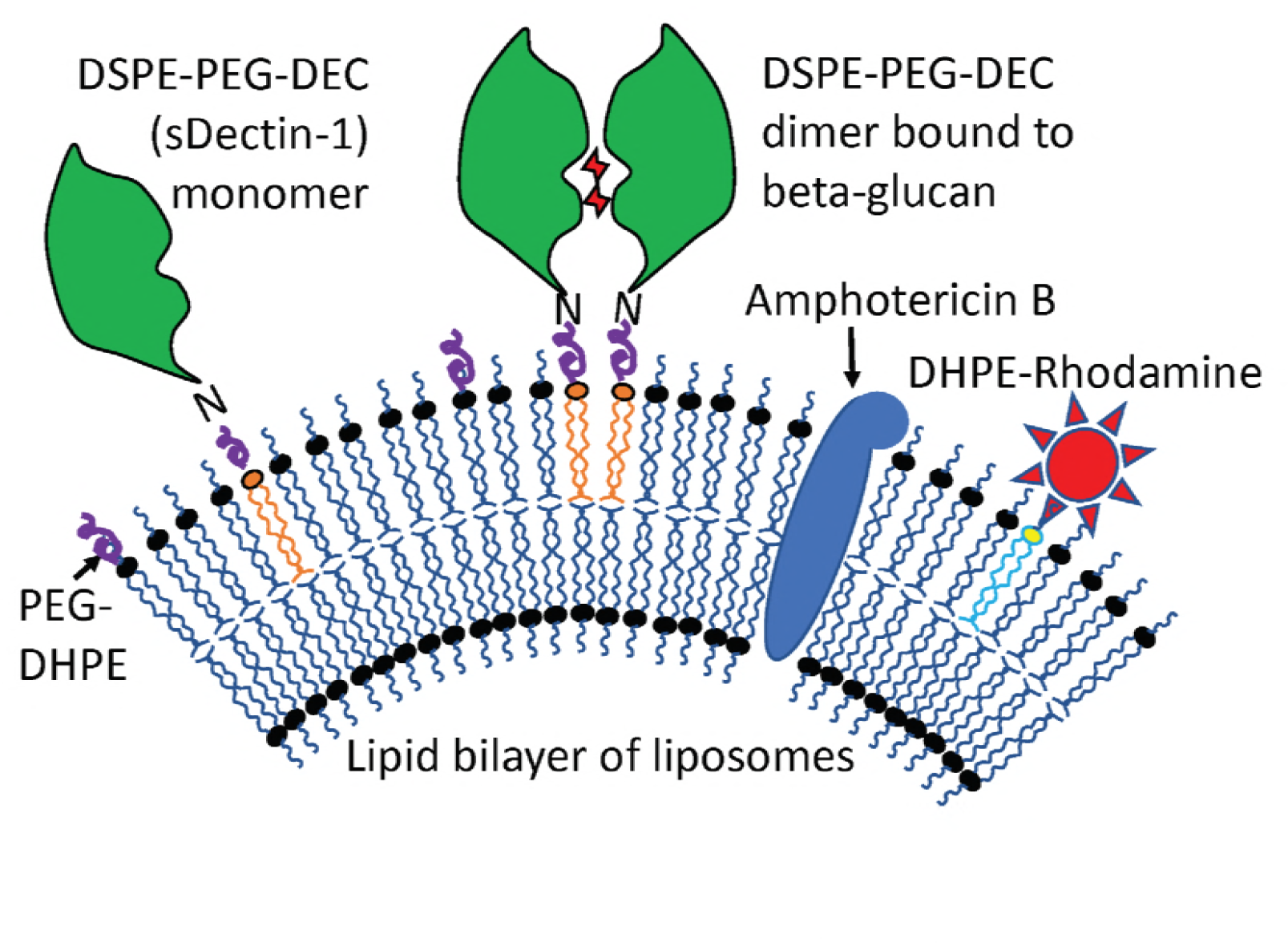
Model of DEC-AmB-LLs, liposomes loaded with sDectin-1, amphotericin B, and rhodamine. Amphotericin B (AmB, blue oval structure) was intercalated into the lipid bilayer of 100 nm diameter liposomes. sDectin-1 (DEC, green globular structure) was coupled to the lipid carrier DSPE-PEG. Both DSPE-PEG-DEC and red fluorescent DHPE-Rhodamine (red star) were also inserted into the liposomal membrane. sDectin-1, Rhodamine, AmB and liposomal lipids were in a 1:2:11:100 mole ratio, respectively (Supplemental Table S1). Two sDectin-1 monomers (two DSEP-PEG-DEC molecules) must float together in the membrane to bind strongly to cell wall beta-glucans (red sugar moieties). The two liposomal controls examined were BSA-AmB-LLs that containing an equal ug amount of 65 kDa BSA in place of 22 kDa sDectin-1 (i.e., 0.33:2:11:100 mole ratio) and AmB-LLs lacking any protein coating (0:2:11:100 mole ratio). From these mole ratios, the surface area of an 100 nm diameter liposome, and the published estimate of 5 × 10^6^ lipid molecules per 10^6^ nm^2^ of lipid bilayer(61), we estimated that there were approximately 3,000 Rhodamine molecules in each liposome preparation and 1,500 sDectin-1 monomers in each DEC-AmB-LL. Note that for simplicity the proper ratios of these molecules are not shown in the figure.

### Targeted liposomes

Liposomes biochemically resemble endogenous exosomes (16–18). They efficiently penetrate the endothelial barrier and reach target cells deep in most major organs for the “passive delivery” of variously loaded therapeutic drugs (19–23). Targeted-liposomes have binding specificity for a plasma membrane antigen to enable the “active delivery” of a packaged therapeutic to diseased cells. Targeting is most commonly achieved with a monoclonal antibody such that immunoliposomes bind a specific cell type or types. Over 100 publications, most focused on particular types of cancer cells, show that targeted immunoliposomes improve the cell-type specificity of drug delivery and reduce toxicity. Therapeutic drug loaded immunoliposomes include those targeting cells expressing the VEGF-Receptors-2 and −3 (24), the oxytocin receptor(25), the epidermal growth factor receptor, EGFR (26), CD4 (27), and HER2 (28, 29). The active delivery of immunoliposomes generally improves cell-type specificity and drug effectiveness by 3- to 10-fold (25, 30, 31) over passive delivery. A wide variety of drugs have been delivered via targeted liposomes including toxins such as doxorubicin, paclitaxel and rapamycin (32, 33), growth hormones such as Transforming Growth Factor-beta (34), and analgesics such as the indomethacin (25). We are unaware of any reports of immunoliposomes specifically targeting antifungals to invasive fungal cells, however, the immuno-targeting of AmB loaded liposomes to the vessel wall of pulmonary capillary cells in *A. fumigatus* infected mouse lungs results in increased mouse survival rates (15). Our experimental hypothesis is that AmB-LLs targeted directly to the cell wall of *A. fumigatus* will have enhanced antifungal activity over current untargeted AmB-LLs.

### Dectin-1 binds beta-glucans on the surface of pathogenic fungi

Dectin-1 is transmembrane receptor expressed in natural killer lymphocytes encoded by the *CLEC7A (C-Type Lectin Domain Containing 7A, beta-Glucan Receptor)* gene in mice and humans. Dectin-1 binds various beta-glucans in fungal cell walls and is the primary receptor for transmembrane signaling of the presence of cell wall components from the surface of fungal cells, stimulating an innate immune response (35–38). Human and mouse Dectin-1 are 244 and 247 amino acid-long plasma membrane proteins, respectively, although there are mRNA splice variants producing shorter human isoforms. Dectin-1 floats in the membrane as a monomer, but binds to beta-glucans as a dimer as modeled in our design of Dectin-1 targeted liposomes shown in **Fig. 1**(39). The 176 amino acid long (20 kDa) extracellular C-terminal, beta-glucan binding domain is often manipulated alone as sDectin-1. The beta 1→3 glucans are a structurally diverse class of polysaccharides, and as such, sDectin-1 binds various beta-glucans differentially with IC50s ranging from 2.6 mM to 2.2 pM (38). sDectin-1 is reported to recognize *A. fumigatus* cell wall components much more efficiently on germinating conidia and germ tubes than on dormant conidia or mature hyphae (40, 41). Having pan-fungal binding activity, Dectin-1 may provide broader antifungal targeting abilities for liposomes than a monoclonal antibody (42).

## Results

### Preparation of amphotericin B loaded sDectin-1 coated liposomes

Pegylated liposomes were remotely loaded with 11 moles percent AmB relative to moles of liposomal lipids to make control AmB-LLs, which are similar in structure and AmB concentration to commercial un-pegylated AmBisomes (**Materials and Methods, Supplemental Table S1**). sDectin-1 (DEC, **Supplemental Fig. S1**, **Fig. S2**) and Bovine serum albumin (BSA) were coupled to a pegylated lipid carrier, DSPE-PEG. One mole percent DSPE-PEG-DEC was incorporated into AmB-LLs to make sDectin-1 coated DEC-AmB-LLs (**Fig. 1**) and 0.33 mole percent DSPE-PEG-BSA was incorporated into AmB-LLs to make BSA-AmB-LLs. This mole ratio of 22 kDa sDectin-1 and 65 kDa BSA results an equivalent ug amounts of protein coating each set of liposomes. Because these protein coated liposomes were made from the same AmB-LLs, all three liposomal preparations contain 11 moles percent AmB relative to moles of lipid. Two moles percent of DHPE-Rhodamine were loaded into all three classes of liposome (**Fig. 1**).

### sDectin-1 coated liposomes DEC-AmB-LLs bind strongly to fungal cells

In assays performed on *A. fumigatus* germlings, rhodamine red fluorescent DEC-AmB-LLs bound strongly to germinating conidia and to germ tubes as shown in **Fig. 2**. The sDectin-1-targeted liposomes often bound in large numbers and in aggregates to particular regions. While 100 nm liposomes are too small to be resolved by light microscopy, individual liposomes are visible as somewhat uniformly sized small red fluorescent dots (orange arrows, **Fig. 2A**), which are easily detected due to their each containing an estimated 3,000 rhodamine molecules (**Fig. 1**). From examinations of larger fields of germlings it appears that essentially all bind DEC-AmB-LLs (**Fig. 2C and 2D**). AmBisome-like AmB-LLs (**Fig. 2B**) and bovine serum albumin coated liposomes, BSA-AmB-LLs (**Fig. 2E & 2F**) did not bind detectably to germinating conidia or germtubes, when tested at the same concentration. Maximum labeling by DEC-AmB-LLs was achieved within 15 to 30 min and the strong red fluorescent signals of DEC-AmB-LLs bound to cells were maintained for weeks, when fixed cells were stored in the dark in PBS at 4°C.

**Fig. 2.**
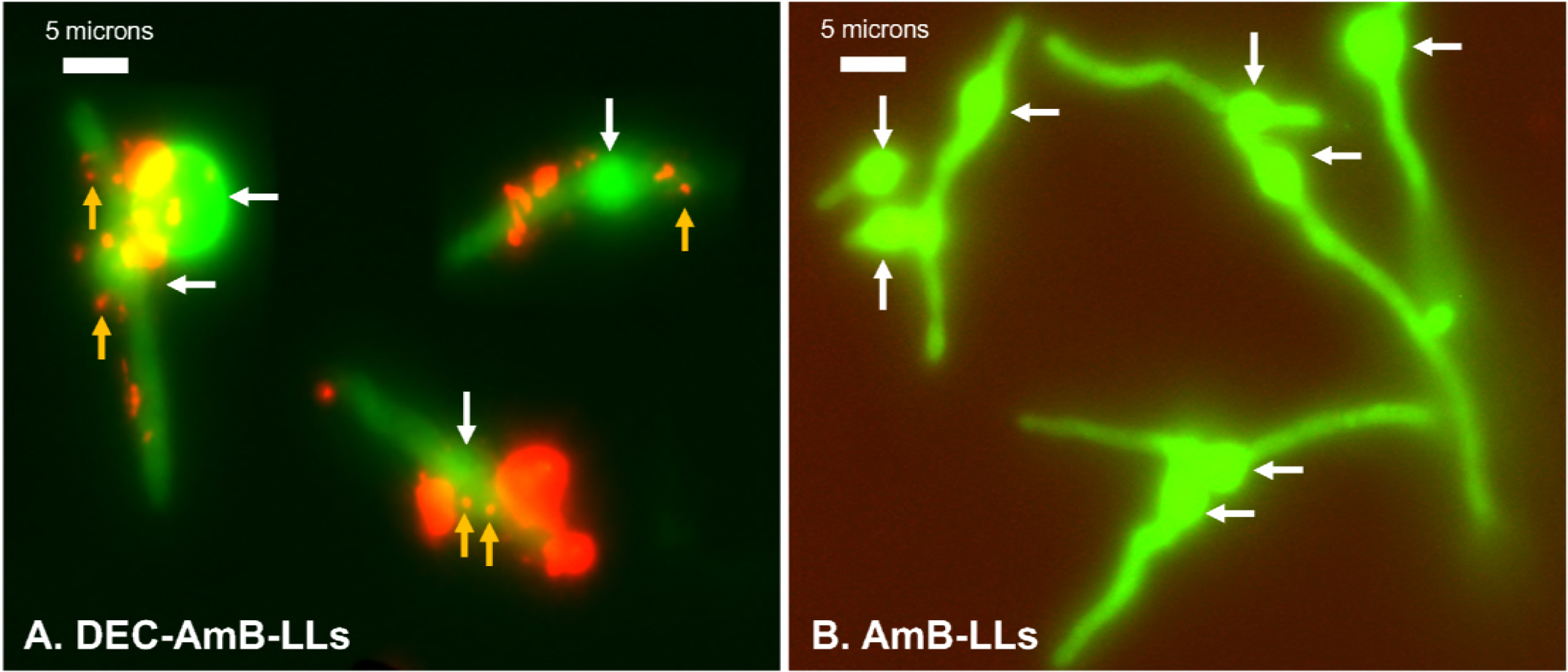
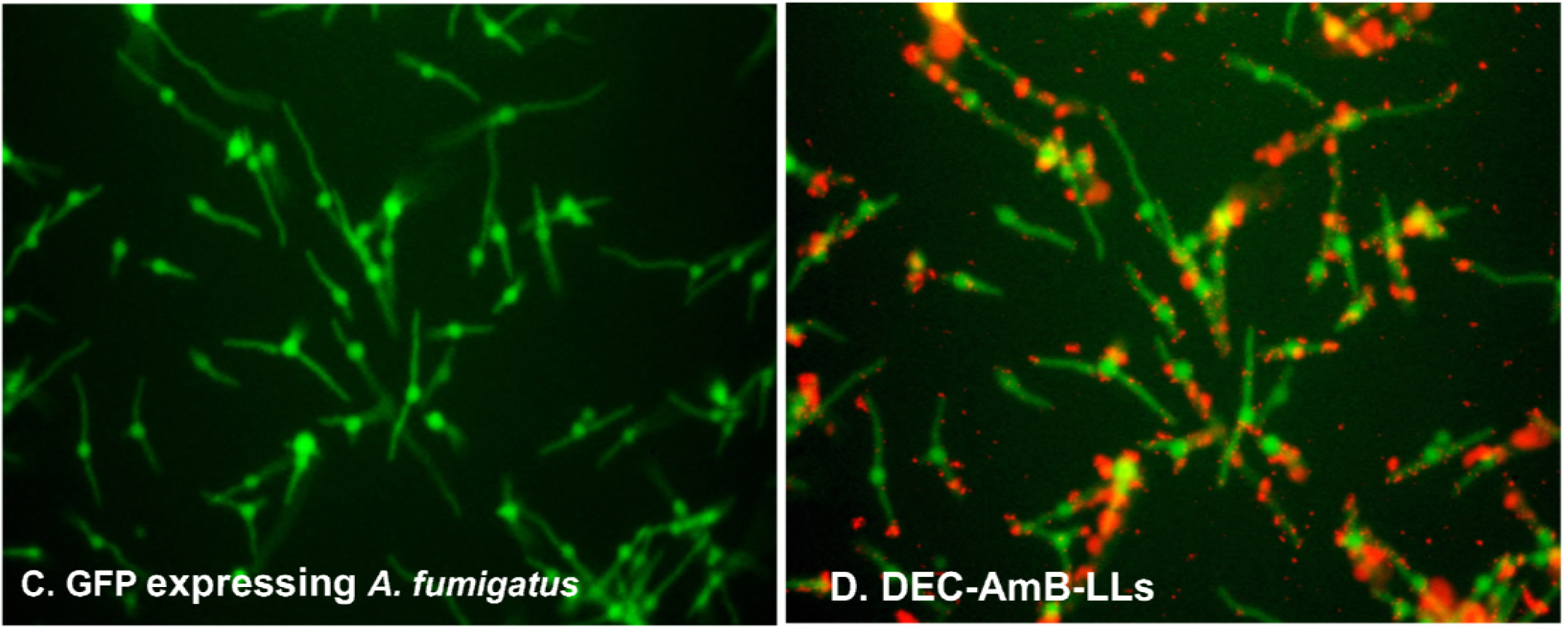
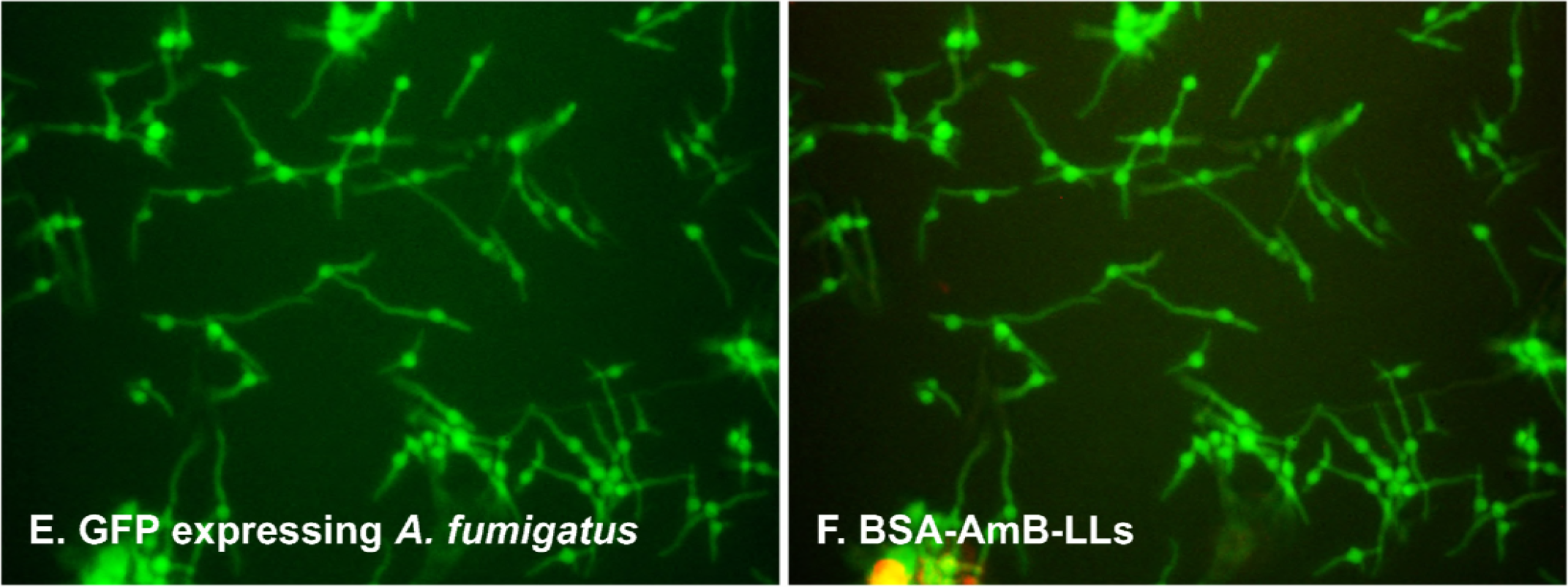
sDectin-1 coated DEC-AmB-LLs bound strongly to germinating conidia and germ tubes of *A. fumigatus*, while AmB-LLs and BSA-AmB-LLs did not. *A. fumigatus* conidia were germinated and grown for 8 to 10 hr in VMM + 1% glucose at 35°C in 24 well microtiter plates before being fixed and stained with fluorescent liposomes. A. Rhodamine red fluorescent DEC-AmB-LLs bound swollen conidia (white arrows) and germ tubes of *A. fumigatus*. B. Rhodamine red fluorescent AmB-LLs did not bind at detectable frequencies. No AmB-LLs were detected even when the red channel was enhanced as in this image. The smallest red dots in plate A represent individual 100 nm diameter liposomes viewed based on their fluorescence (orange arrows). Large clusters of liposomes form the more brightly red stained areas. C and D were stained with DEC-AmB-LLs. E and F stained with BSA-AmB-LLs. A through F. Cells were grown for 8 to 10 hr in VMM + 1% glucose at 37°C. Labeling was performed in LDB for 60 min. All three liposomes preparations were diluted 1:100 such that liposomal sDectin-1 and BSA proteins were at final concentrations of 1 ug/100 uL. Germlings were viewed in the green channel alone for cytoplasmic fluorescent EGFP expression and red channel for rhodamine fluorescent liposomes. A and B were photographed at 63X under oil immersion in a compound fluorescence microscope and red florescence was further enhanced in B to detect potentially individual liposomes. C through F were photographed at 20x on an inverted fluorescence microscope.

DEC-AmB-LLs also bound to germinating conidia and most hyphae from more mature cultures as shown in **Fig. 3**. Again, the sDectin-1-targeted liposomes often bound in aggregates, but some fairly uniformly sized individual small red dots are visible (orange arrows, **Fig. 3A**), which appear to be individual fluorescent liposomes. AmB-LLs did not bind significantly to older conidia or mature hyphae (**Fig. 3E & 3F**) nor did BSA-AmB-LLs.

**Fig. 3.**
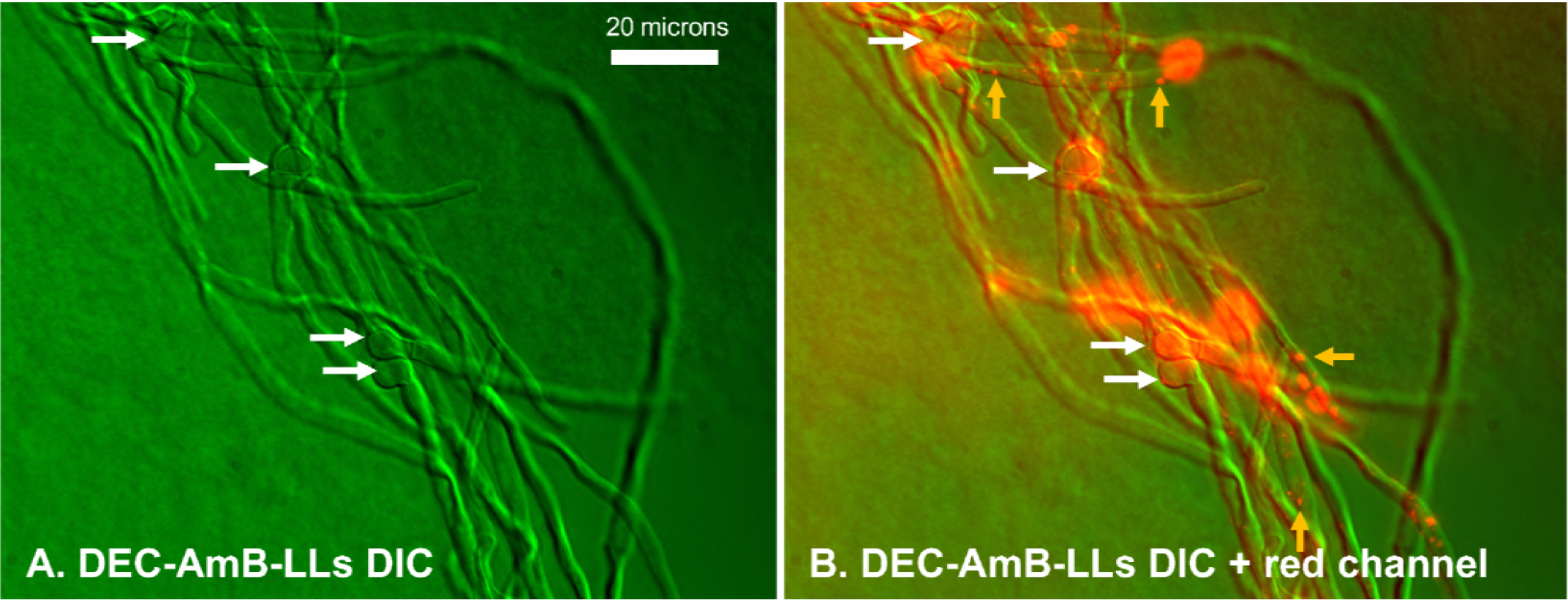
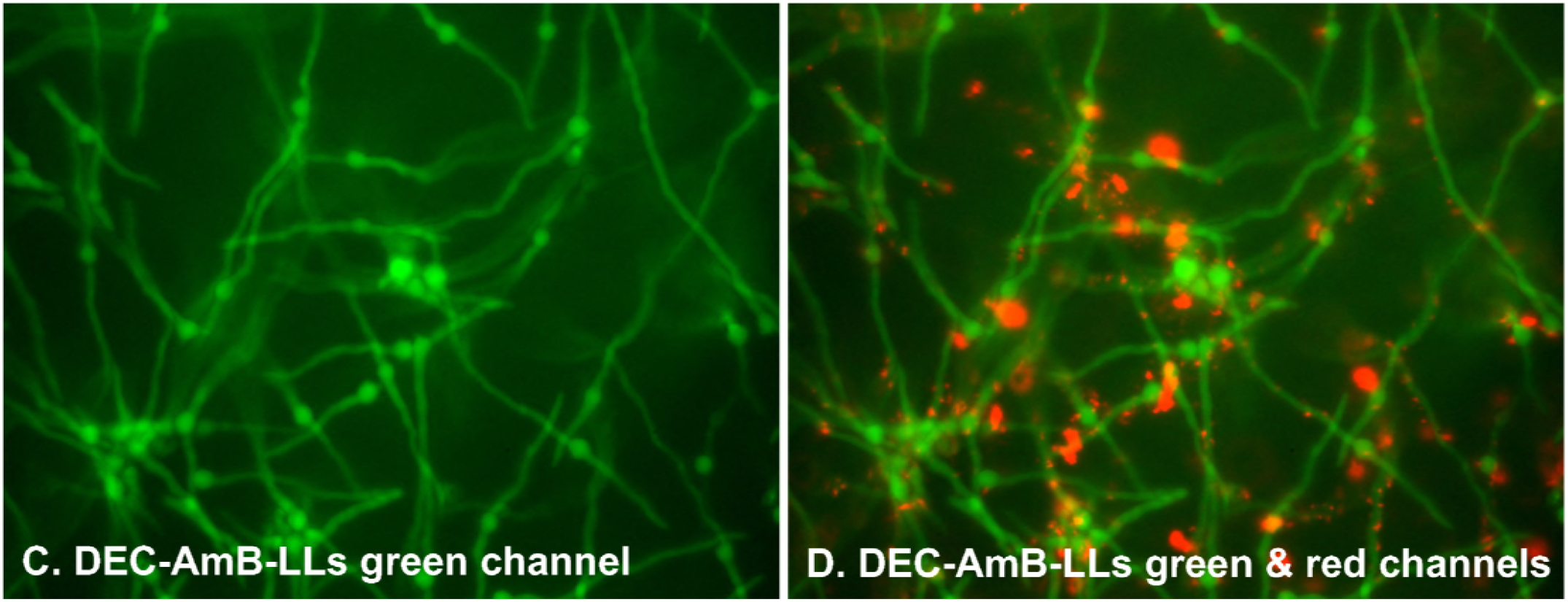
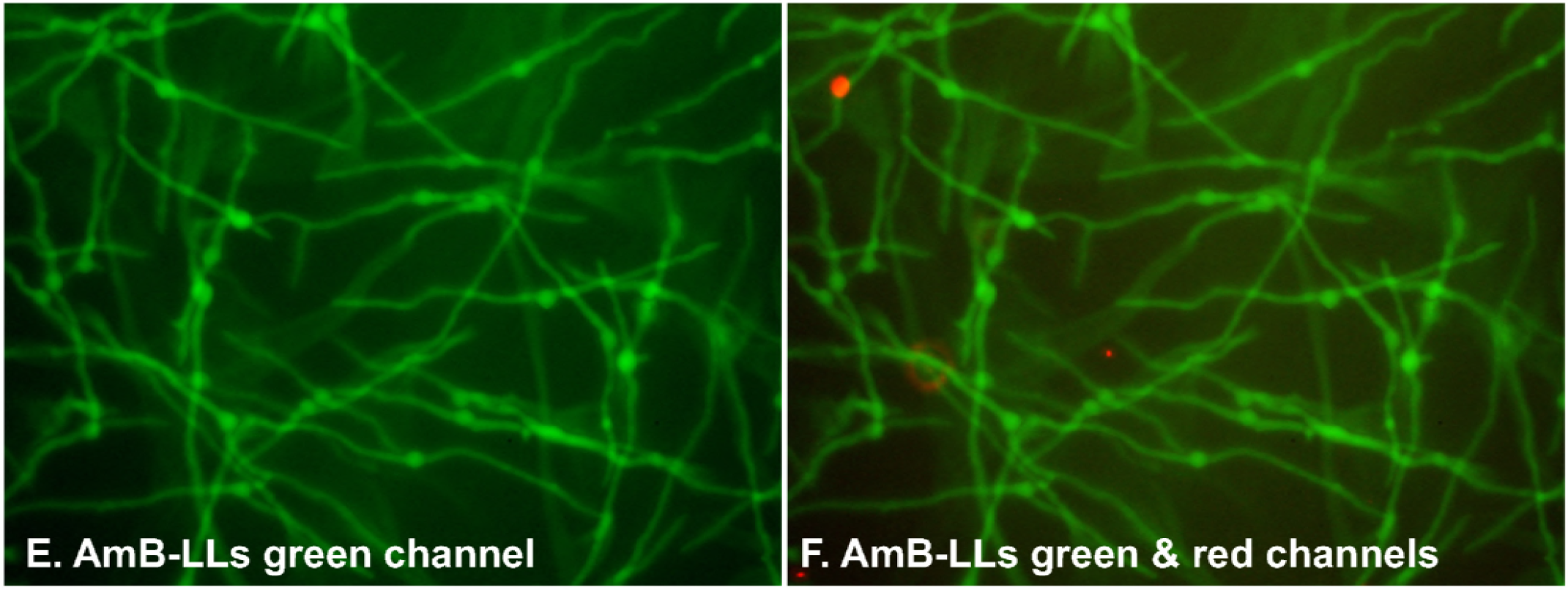
sDectin-1 coated DEC-AmB-LLs bound germinating conidia and hyphae of mature *A. fumigatus* cells, while untargeted AmB-LLs and BSA-AmB-LLs did not. *A. fumigatus* conidia were germinated and grown for 16 hr in VMM + 1% glucose at 35°C in 24 well microtiter plates before being fixed and stained with fluorescent liposomes. A. through D. Cells were stained with rhodamine red fluorescent DEC-AmB-LL diluted 1:100 such that sDectin-1 was at 1 ug/100 uL, and E. and F. with the equivalent amount of red fluorescent AmB-LLs for 60 min. A. DIC image alone. B. Combined DIC and red fluorescence image. A and B. show that Rhodamine fluorescent DEC-AmB-LLs bound to germinating conidia (white arrows) and hyphae. In B the smallest red dots represent individual 100 nm liposomes (orange arrows). C through F examined cytoplasmic green fluorescent EGFP and the red fluorescence of liposomes. C and D show that nearly all conidia and most hyphae stained with DEC-AmB-LLs. E & F show that AmB-LLs did not bind. A and B were photographed at 63X under oil immersion and C through F were photographed at 20X on an inverted fluorescent microscope.

On plates covered with dense layers of mature hyphae, the number of bound liposomes and liposome aggregates were counted in multiple fluorescent images. DEC-AmB-LLs bound to both formalin fixed (**Fig. 4A-C**) and live (**Fig. 4D-F**) *A. fumigatus* cells 100- to 200-fold more efficiently than AmB-LLs or BSA-AmB-LLs. Labeling by DEC-AmB-LLs was inhibited 50-fold by the inclusion of soluble beta-glucan, laminarin, but not sucrose, confirming that binding was beta-glucan specific (**Fig. 4G-4GI**). Finally, DEC-AmB-LLs labeled *Cryptococcus neoformans* cells and *Candida albicans* pseudohyphae (**Supplemental Fig. S3**), while control liposomes did not. In short, Dectin-coated Amphotericin B loaded liposomes bound efficiently to a variety of fungal cells, while control liposomes did not.

**Fig. 4.**
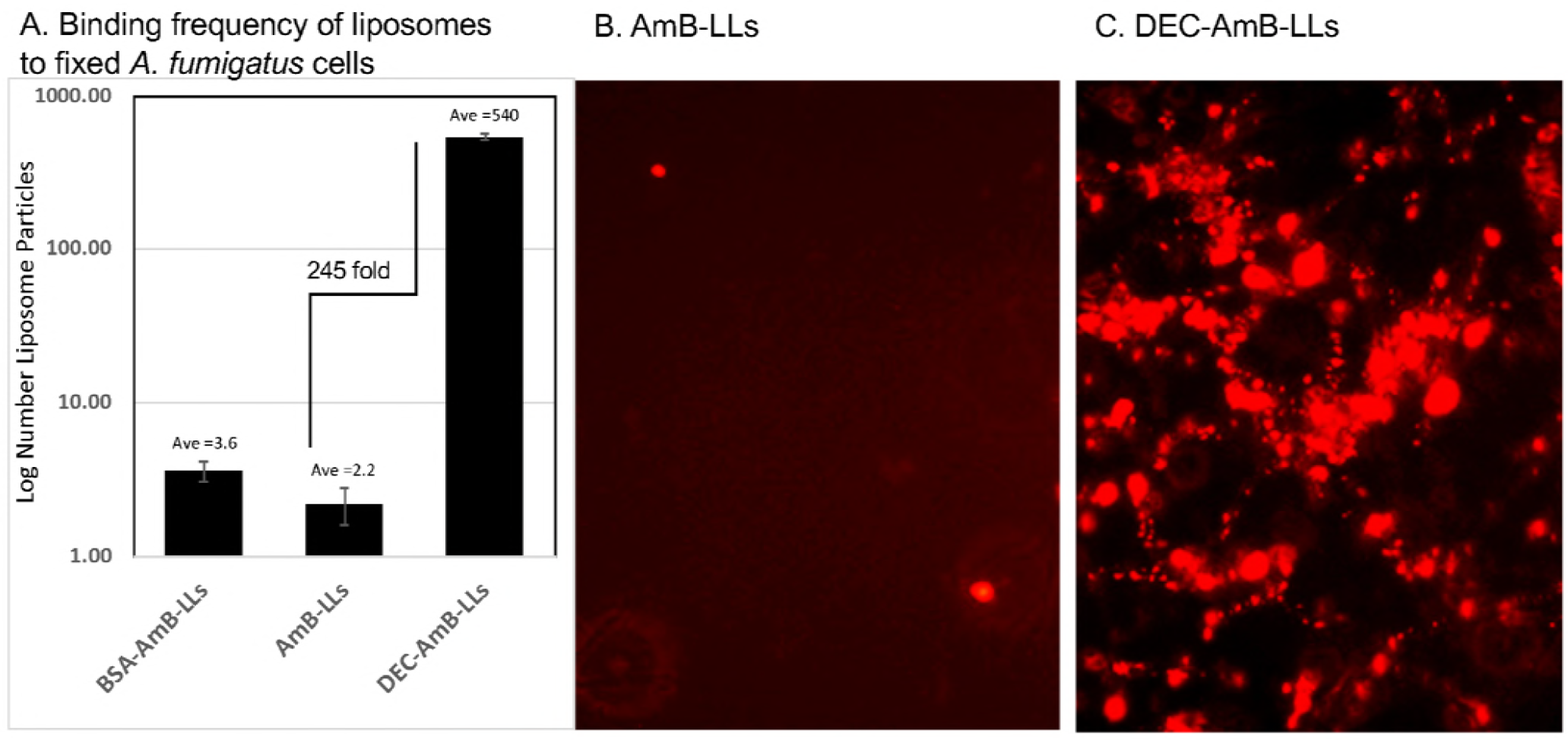
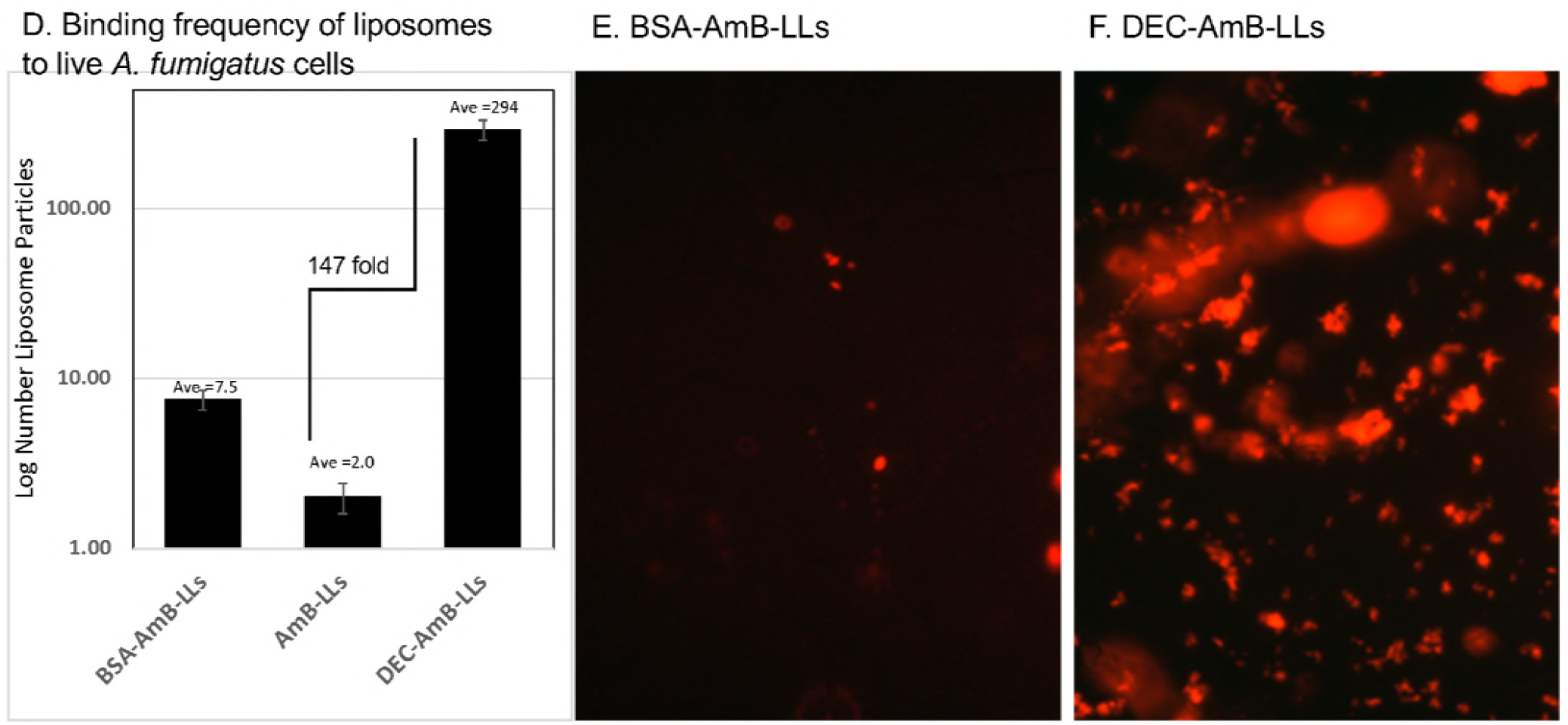
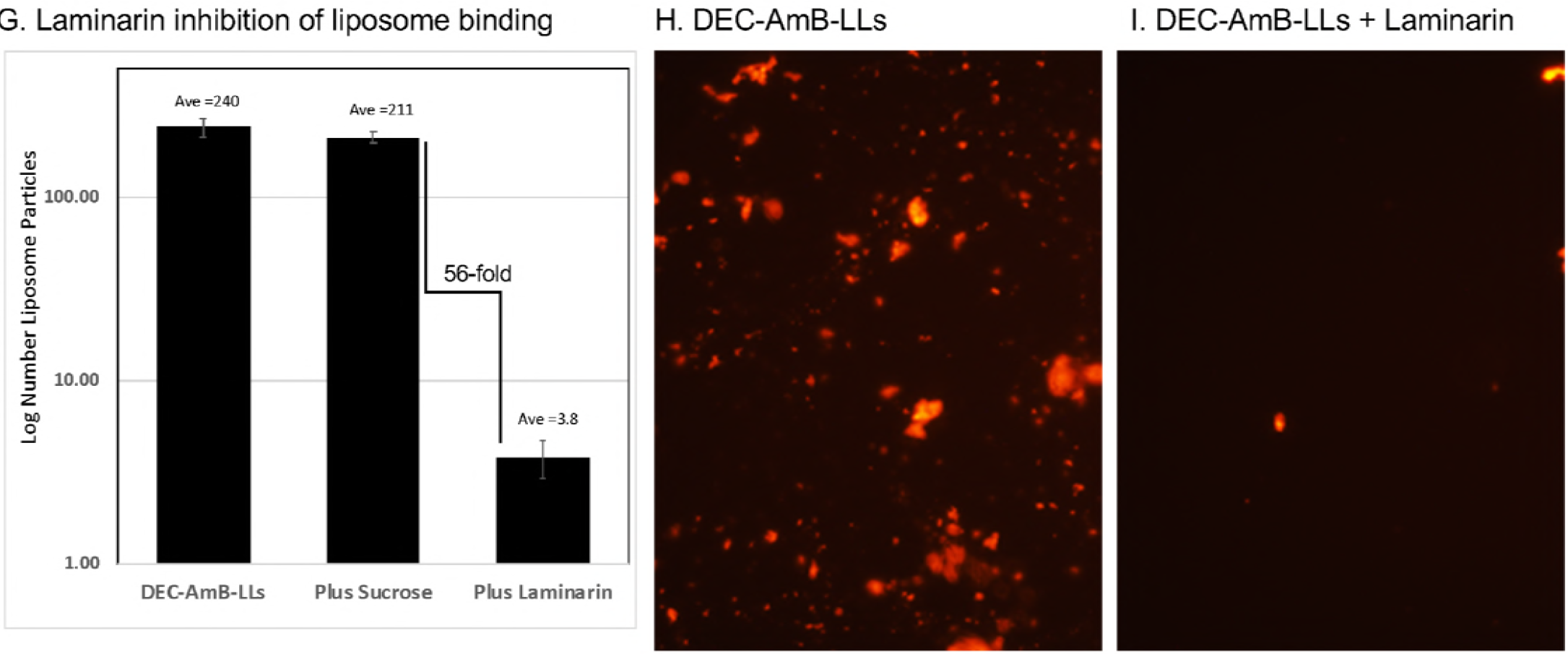
sDectin-1-coated DEC-AmB-LLs bound two orders of magnitude more frequently to *A. fumigatus* than control AmB-LLs and binding was inhibited by a soluble beta-glucan. Samples of 4,500 *A. fumigatus* conidia were germinated & grown at 35°C for 36 hours VMM+1% glucose, fixed in formalin or examined live, and incubated for 1 hr with 1:50 dilutions of liposomes in liposome dilution buffer. Unbound liposomes were washed out. Multiple fields of red fluorescent images were photographed at 20X and red fluorescence enhanced equivalently for all images. Each photographic field contained approximately 25 swollen conidia and an extensive network of hyphae (not shown). A, B, C. Labeling formalin fixed cells. D, E, F. Labeling live cells. G, H, I. Inhibition of DEC-AmB-LL labeling of fixed cells by 1 mg/mL laminarin, a soluble beta-glucan vs 1 mg/mL sucrose as a control. A, D, & G. The number of red fluorescent liposomes and clusters of liposomes were counted, averaged per field and plotted on a log_10_ scale. The numerical average is indicated above each bar and on the vertical axis. Standard errors are shown. Examples of photographic fields of liposomes used to construct the adjacent bar graphs are shown in B, C, E, F, H, and I.

### Killing and growth inhibition of fungi by DEC-AmB-LLs

We performed various fungal cell growth and viability assays, after treating *A. fumigatus* with liposomes delivering AmB concentrations near its estimated ED50 of 2 to 3 uM AmB (43) or below its estimated MIC of 0.5 uM for various strains of *A. fumigatus* (44). In most of these experiments, 4,500 conidia were germinated and incubated for 12 to 72 hr in 96 well microtiter plates along with drug loaded liposomes. Longer incubation times were often needed to resolve differences among sthe liposome preparation delivering higher concentrations of AmB. **Fig. 5** shows that targeted DEC-AmB-LLs killed or inhibited the growth of *A. fumigatus* cells far more efficiently than BSA-AmB-LLs or uncoated AmB-LLs delivering the same concentrations of AmB. Assays with CellTiter-Blue reagent, which assesses cytoplasmic reductase activity as a proxy for cell integrity and viability, showed that treating cells with DEC-AmB-LLs delivering 3 uM AmB killed *A. fumigatus* more than an order of magnitude more effectively than AmBisome-like AmB-LLs or BSA coated liposomes BSA-AmB-LL (**Fig. 5A**). As a second method to score liposomal AmB activity, we measured hyphal length. Hyphal length assays gave a similar result, showing that DEC-AmB-LLs delivering 3 uM AmB were far more effective at inhibiting hyphal growth than AmB-LLs or BSA-AmB-LLs (**Fig 5B**). In a complete biological replicate experiment with an independient slightly different method of preparaing liposomes we obtained a similar although less dramatic result, when delivering 3 uM AmB (**Fig. 5C and 5D**).

**Fig. 5.**
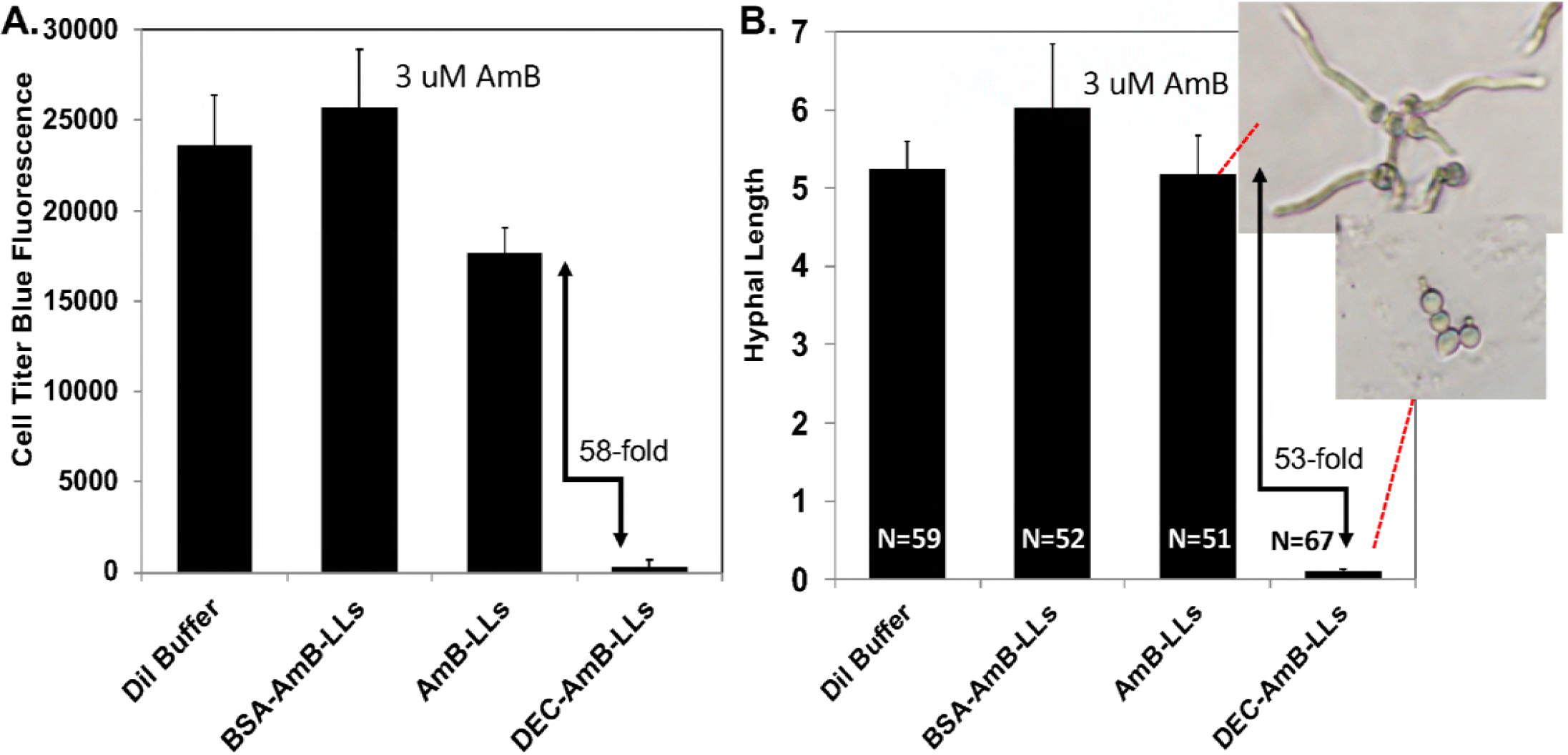
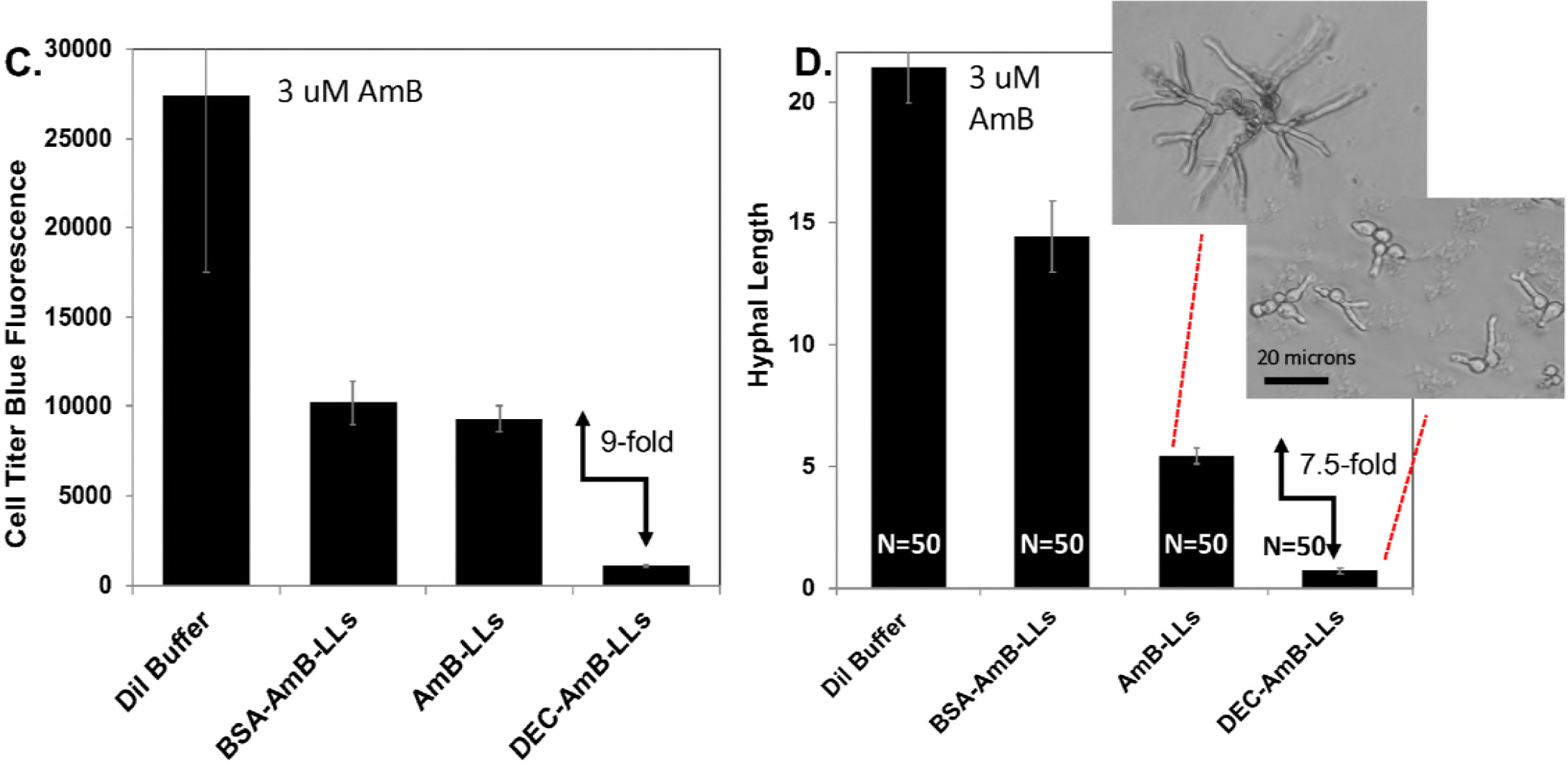
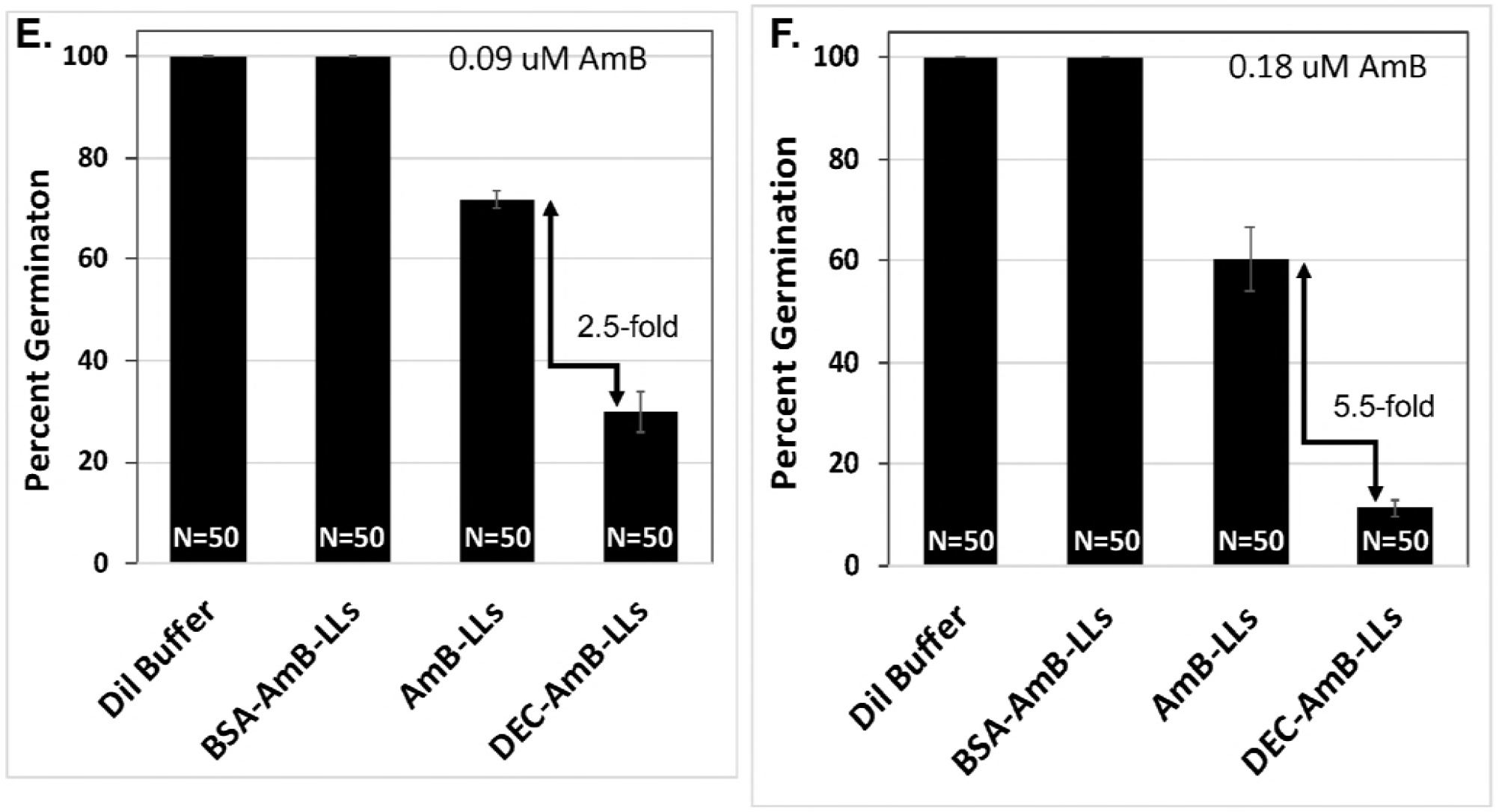
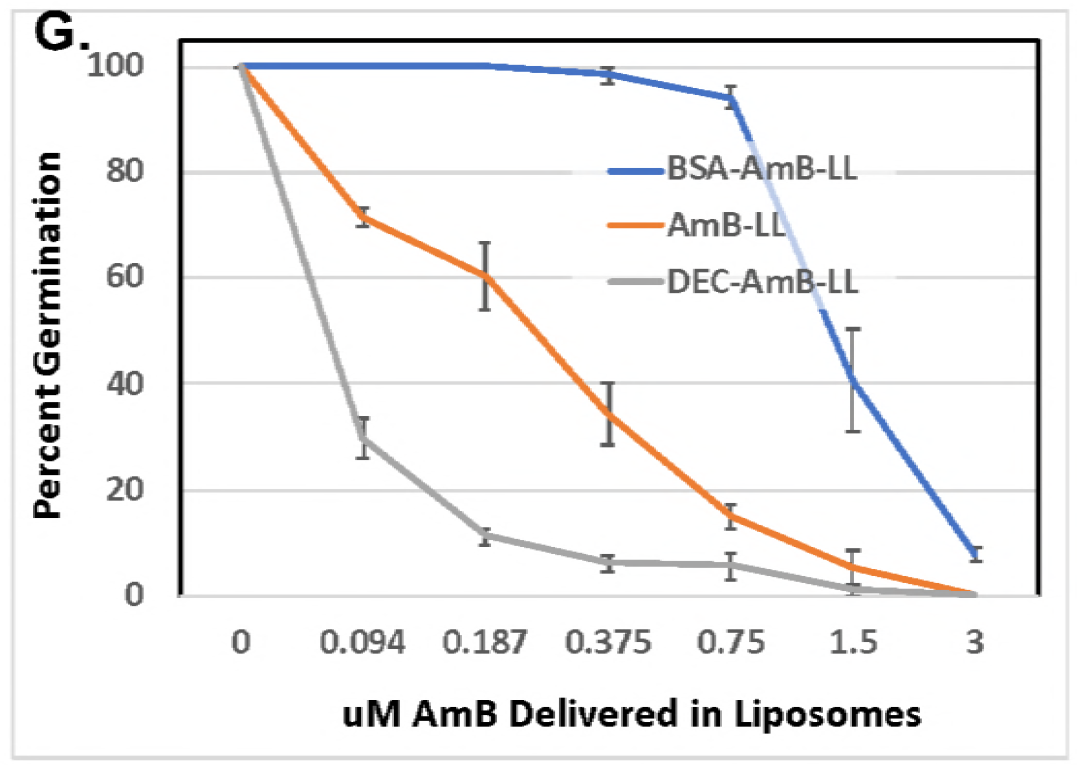
DEC-AmB-LLs inhibited the growth *A. fumigatus* far more efficiently than AmB-LLs. Samples of 4,500 *A. fumigatus* conidia were germinated & grown in 96 well microtiter plates in Vogel’s Minimal Media (VMM+1% glucose) for 8 to 56 hr at 35°C and treated at the same time with liposome preparations delivering the indicated concentrations of AmB to the growth media (A-D 3 uM AmB, a 1:300 fold dilution of all three liposome preparations), E. 0.09 uM, F. 0.18 uM, G. 0.9 to 3 uM) or an equivalent amount of liposome dilution buffer. Viability and growth were estimated using CellTiter-Blue reagent (A and C) or by measuring hyphal length (B & D) or by scoring percent germination (E, F, and G). Background fluorescence from wells with CellTiter-Blue reagent in the media, but lacking cells and liposomes was subtracted. Std. Errors are indicated. Inset photos in B and D show examples of the length of hyphae assayed for AmB-LLs and DEC-AmB-LL treated sample. One unit of hyphal length in B and D equals 5 microns. Cells were grown for 8 to 56 hrs. A and B and C and D compare the results from two biological replicate experiments with independently conjugated sDectin-1 and assembled liposomes.

A third assay of liposomal AmB activity was employed, which measured the percent of conidia that germinated in the presence of the various liposomal preparations (**Fig. 5E and 5F**). DEC-AmB-LLs delivering as little as 0.09 uM and 0.187 uM AmB inhibited the germination of *A. fumigatus* conidia significantly better than AmB-LLs or BSA-AmB-LL. The dose response to DEC-AmB-LL was based on a germination assay performed after a fixed period of growth using AmB concentrations from 0.09 to 3 uM is shown in **Fig. 5G** DEC-AmB-LLs outperform AmB-LLs and BSA-AmB-LLs over a wide range of concentrations.

### Reduced animal cell toxicity of DEC-AmB-LLs

AmB-LLs and AmB deoxycholate micelles were slightly more toxic to HEK293 human embryonic kidney cells than DEC-AmB-LLs or BSA-AmB-LLs based on CellTiter-Blue assays of cell viability (**Supplemental Fig. S4**).

## Discussion

We demonstrated that sDectin-1 targeted DEC-AmB-LLs are significantly more effective at binding to and inhibiting the growth of fungal cells and are slightly less toxic to human cells than uncoated AmB-LLs. The biochemical manipulation of mouse or human sDectin-1 has been complicated, because the proteins easily aggregate and become insoluble and inactive in aqueous buffers. This problem is perhaps in part, because they are composed of 11 to 13% hydrophobic amino acids and contain three disulfide crosslinks in their carbohydrate recognition domains. A wide variety of protein chemical manipulations have been applied to improve sDectin-1 solubility with mixed success. For example, the solubility and utility of sDectin-1 was increased by tethering it to the 56 amino acid long B1 domain of Streptococcal Protein G (45) or more commonly to the 232 a.a. long Fc constant region of IgG1 antibody (41). Bacterially produced non-glycosylated sDectin-1 renatured from inclusion bodies and de-glycosylated and native mammalian cell produced sDectin-1 all retain indistinguishable beta-glucan binding activity (39, 46, 47). Therefore, we proceeded with sDectin-1 production in *E. coli*, with the potential for highest yield and lowest cost. We overcame sDectin-1’s solubility problems by combining a variety of old and new approaches including, (1) the use of a very short charged peptide tag, (2) the inclusion of 6 M guanidine hydrochloride (GuHCl) during protein extraction, purification, and chemical modification, (3) the inclusion of the protein solubilizing agent, arginine, during renaturation, liposomal loading, and storage, and (4) the inclusion of a sulfhydryl reducing agent BME in all steps.

sDectin-1 is reported to bind efficiently to *A. fumigatus* germinating conidia and germ tubes, but inefficiently if at all to mature hyphae and not at all to un-germinated conidia (40, 41, 48). Our data with sDectin-1 coated AmB loaded liposomes are partially consistent with previous observations, except that we observed reasonable efficient staining of mature hyphae (**Fig. 3 & 4)**. Poor cell or hyphal binding may be explained by polymorphic expression of beta-glucans in different stages of fungal cell growth (48, 49). During infection, however, interaction with host immune cells reportedly expose otherwise masked β-glucans (50), enhancing the potency of the targeted DEC-AmB-LLs. Herein, sDectin-1 coated fluorescent DEC-AmB-LLs bound efficiently to germinating conidia, germ tubes, and hyphae, suggesting our modified sDectin-1 presented on the surface of liposomes retained its ability to form complexes with affinity for fungal beta-glucans expressed at various stages of growth. Our data showed for the first time that in vitro chemically modified sDectin-1 (DSPE-PEG-DEC) retained its fungal cell binding specificity. Furthermore, DEC-AmB-LLs binding was rapid and remained stably bound to cells for weeks. Perhaps the greater avidity of liposome coated with ~1,500 sDectin-1 molecules insured the rapid efficient binding and very slow release of bound liposomes, parallel to the avidity of pentameric IgM antibody. The presence of thousands rhodamine molecules on each liposome (**Fig. 1**) should have greatly increased the chance of detecting unambiguous fluorescent signals relative to detecting the binding of sDectin-1 dimers as reported in previous studies.

In a large number of experiments using different binding buffers including BSA blocker and various incubation periods we never detected any significant affinity of uncoated AmB-LLs or BSA-AmB-LLs for fungal cells, with one exception. In preliminary experiments in which BSA blocker was omitted during the incubation, we observed BSA-AmB-LLs bound modestly well to *A. fumigatus* germinated conidia, while we still did not observe AmB-LLs binding. By contrast, Chavan et al (51) detected efficient binding of fluorescent pegylated liposomes to primary tips and septa on *A. fumigatus* hyphae even in the presence of serum (51). We cannot account for this disparity between their data and ours, except that their liposomes had a different lipid composition and lacked both sDectin-1 and amphotericin B (**Supplemental Table S1**).

*Aspergillus, Candida* and *Cryptococcus species* belong to three evolutionarily disparate groups of fungi, the Hemiascomycetes, Euascomycetes, and Hymenomycetes, respectively, which are separated from common ancestry by hundreds of millions of years (52). DEC-AmB-LLs bound specifically to all three. This suggests the beta-glucans found in the outer cell wall of many pathogenic fungi will be conserved enough in structure and accessible enough bind sDectin-1 targeted liposomes.

In various biological and experimental replicate experiments using different assay methods we showed that DEC-AmB-LLs killed or inhibited *A. fumigatus* cells far more efficiently than AmBisome-like AmB-LLs delivering the same level of AmB. In all of our experiments, DEC-AmB-LLs were from several fold to more than an order of magnitude more fungicidal than control liposomes over a wide variety of AmB concentrations tested that were near or below the estimated ED50 of 3 uM. We detected significantly stronger activity of DEC-AmB-LLs over

AmB-LLs even at AmB concentrations as low as 0.094 uM AmB, well below AmB’s MIC. DEC-AmB-LLs significantly decreased the amount of AmB required for an ED50 or a MIC for *A. fumigatus*. The time of incubation with drug loaded liposomes strongly influences the ability to resolve differences among the three liposome preparations. For example, when all three liposome preparations delivered high AmB concentrations (e.g., 0.75, 1.5 and 3 uM) they caused a lag in the germination of conidia. Hence, longer incubation periods were needed to allow sufficient fungal growth to resolve the improved performance of DEC-AmB-LLs. Short incubation periods were needed to resolve differences at low AmB concentrations (e.g., 0.37, 0.18, 0.94 uM). Thus we were unable to obtain a dose response curve that reflected the optimal performance of DEC-AmB-LLs over a wide range of AmB concentrations.

Looking forward, there are a number of important variables we have not yet explored. For example, we coated liposomes with a single concentration of sDectin-1, approximately 1,500 molecules per liposome and do not know if this is the optimal concentration for cell binding and drug delivery. Also, although DEC-AmB-LL were superior in all aspects to AmB-LLs we do not know the ratio of growth inhibition to killing at different AmB concentrations. Efficient killing can be followed by rapid outgrowth of the remaining cells obscuring the results. This is particularly relevant to the treatment of aspergillosis, because current drug formulations only partially reduce the fungal cell load in mice and humans. Finally, an important next step in our research needs be an examination of the performance sDectin-1 coated antifungal drug loaded liposomes in animal models of fungal diseases.

In summary, sDectin-1 conjugated to a pegylated lipid carrier and inserted as monomers into liposomes must float together to form functional dimers or multimers as they bind beta-glucans or we would not have observed the strong efficient binding of DEC-AmB-LLs to fungal cells. Our DEC-AmB-LLs efficiently bind beta-glucans in the cell walls of diverse fungal species. Multiple growth and viability assays on DEC-AmB-LLs delivering AmB concentrations from 0.094 to 3 uM suggest that sDectin-1 coated liposomes greatly improved the performance of liposomal AmB. Taking these results altogether, it is reasonable to propose that sDectin-1 coated liposomes have significant potential as pan-fungal carriers for targeting antifungal therapeutics.

## Materials and Methods

### Fungal growth

*Aspergillus fumigatus* strain A1163 was transformed with plasmid pBV126 described in Kang et al. (53) carrying green fluorescent protein EGFP under the control of *Magnaporthe oryzae* ribosomal protein 27 promoter to make strain AEK012. AEK012 was used to monitor fungal cells in some experiments. *A. fumigatus* spores were inoculated on poly-L-lysine coated plates containing Vogel’s Minimal Media (VMM, 1% glucose, 1.5% Agar) and grown for 7 days, at which time conidia were collected in PBS + 0.1% Tween. For fluorescent liposome localization and for growth inhibition and killing assays 20,000 and 4,500 AEK012 conidia were plated on 24 well and 96 well plates, respectively, in VMM, 1% glucose, 0.5% BSA at 35°C for various time periods ranging from 8 hr to 56 hr (54, 55). *Candida albicans* Sc5314 and *Cryptococcus neoformans* H99 were pre-grown in YPD liquid media for overnight. The cells were then washed 3 times with sterile water and resuspended in VMM media and grown on poly-L-lysine coated plates at 35°C for 10 hours. All fungal cell growth was carried out in a BSL2 laboratory. Prior to liposomal staining most fungal preparations were washed 3 times with PBS, fixed in 4% formaldehyde in PBS for 15 to 60 mins, washed twice and stored at 4°C in PBS.

### Production of soluble Dectin-1

The sequence of the codon-optimized *E. coli* expression construct with MmsDectin-1lyshis (NCBI BankIT #2173810) cloned into pET-45B (GenScript) is shown in **Supplemental Fig. S1**. The sequence encodes a slightly modified 198 a.a. long sDectin-1 protein containing a vector specified N-terminal (His)_6_ affinity tag, a flexible GlySer spacer, two lysine residues, another flexible spacer followed by the C-terminal 176 a.a. long murine sDectin-1 domain. *E. coli* strain BL21 containing the MmsDectin-1-pET45B plasmid were grown overnight in 1 L of Luria broth without IPTG induction (**Supplemental Fig. S2**). Modified sDectin-1 was extracted from cell pellets in pH = 8.0, 6 M GuHCl (Fisher BioReagents BP178), 0.1 M Na2HPO4/NaH2PO4, 10 mM Triethanolamine, 100 mM NaCl, 5 mM BME, 0.1% Triton-X100, which was modified from a GuHCl buffer used an earlier study (56). sDectin-1 was bound to a nickel affinity resin (QiaGen, #30210) in this same buffer, washed in same adjusted to pH 6.3, and eluted in this buffer adjusted to pH 4.5. The pH of the eluted protein was immediately neutralized to pH 7.2 with 1 M pH 10.0 M triethanolamine for long term storage. Forty mg of greater than 95% pure protein was recovered per liter of Luria broth (**Supplemental Fig. S2**). Samples of sDectin-1 at 6 ug/uL in this same GuHCl buffer were adjusted to pH 8.3 with 1 M pH =10 triethanolamine and reacted with a 4-molar excess of DSPE-PEG-3400-NHS (Nanosoft polymers, 1544-3400) for 1 hr at 23°C to make DSPE-PEG-DEC. Gel exclusion chromatography on Bio-Gel P-6 acrylamide resin (Bio-Rad #150-0740) in renaturation and storage buffer RN#5 (0.1 M NaH2PO4, 10 mM Triethanolamine, pH 8.0, 1 M L-Arginine, 100 mM NaCl, 5 mM EDTA, 5 mM BME) removed un-incorporated DSPE-PEG and GuHCl (45, 57). DSPE-PEG-BSA was prepared from bovine serum albumin BSA (Sigma, A-8022) by the same protocol, minus the GuHCl from DSPE-PEG labeling buffers and L-Arginine from RN#5 buffer.

### Remote loading of Amphotericin B, sDectin-1, BSA, and Rhodamine into liposomes

Sterile pegylated liposomes were obtained from FormuMax Sci. Inc. (DSPC:CHOL:mPEG2000-DSPE, 53:47:5 mole ratio, 100 nm diameter, 60 umole/mL lipid in a liposomal suspension, FormuMax #F10203A). Small batches of liposomes were remotely loaded with 11 moles percent Amphotericin B (AmB, Sigma A4888) relative to 100% liposomal lipid to make AmBisome-like AmB-LLs used throughout this study. For example, AmB (2.8 mg, 3 umoles, 20 moles %) AmB was dissolved in 13 uL DMSO by heating 10 to 20 min and with occasional mixing at 60°C to make an oil-like clear brown AmB solution. Two hundred and fifty uL of sterile liposomal suspension (15 umoles of liposomal lipid in 50% liposome suspension) was added to the AmB-oil and mixed on a rotating platform for 3 days at 37°C, followed by centrifugation for 10min at 100xg to pellet the AmB oil. Dissolving this oil phase in 0.5 mL DMSO and spectrophotometry at A407 relative to AmB standards in DMSO showed that 1.3 umoles AmB remained undissolved and 1.7 umoles of AmB (11 moles percent) were retained in liposomes. Subsequent gel exclusion chromatography of loaded liposomes over a 10 mL BioGel A-0.5 M agarose resin (BioRad 151-0140) revealed no detectable AmB in the salt volume and essentially all of the AmB was retained by liposomes. Longer incubations resulted in higher percentages of AmB in liposomes. AmB Commercial AmBisomes^®^ (Gilead) are not pegylated and contain 10.6 moles percent AmB relative to lipid (**Supplemental Table S1**).

The DSPE-PEG-sDectin-1 and DSPE-PEG-BSA conjugates in RN#5 buffer and PBS, respectively, were integrated via their DSPE moiety into the phospholipid bilayer membrane of AmB-LLs at 1.0 and 0.33 moles percent of protein relative to moles of liposomal lipid by 60 min incubation at 60°C to make DEC-AmB-LLs and BSA-AmB-LLs. During this same 60°C incubation, the red fluorescent tag, Lissamine rhodamine B-DHPE (Invitrogen, #L1392) was also incorporated at two moles percent relative to liposomal lipid (58–60). Gel exclusion chromatography on BioGel A-0.5 M resin confirmed that Rhodamine and protein insertion into liposomes were essentially quantitative. DEC-AmB-LLs stored at 4°C in RN#5 retained fungal cell binding specificity for 2 months.

### Microscopy of liposome binding

Formalin fixed or live fungal cells were incubated with liposomes in liposome dilution buffer LDB (PBS pH 7.2, 5% BSA, 1 mM BME, 5 mM EDTA) and unbound liposomes washed out after 15 min to 2 hr incubation. Images of rhodamine red fluorescent liposomes, green EGFP *A. fumigatus* and differential interference contrast (DIC) illuminated cells were taken on microscope slides under oil immersion at 63x on a Leica DM6000B automated microscope. Five to six Z-stack images were recorded at one micron intervals and merged in Adobe Photoshop CC2018 using the Linear Dodge method. Bright field and red and green fluorescent images were taken directly of cells on microtiter plates at 20X and 40X on an Olympus IX70 Inverted microscope and an Olympus PEN E-PL7 digital camera and the bight field and/or colored layers merged in Photoshop.

### Cell growth and viability assays

Liposomal stocks were stored at 900 uM AmB and diluted first 5 to 10-fold into liposome dilution buffer LDB with 0.5% BSA and then into VMM or LDB with 0.5% BSA for use at the indicated final AmB concentrations. CellTiter-Blue cell viability assays were conducted as per the manufacturer’s instructions (Promega, document #G8080) using 20 uL of resazurin reagent to treat 100 uL of fungal or animal cells in growth media and incubating for 2 to 4 hours at 37°C. Red fluorescence of electrochemically reduced resorufin product (Ex485/Em590) was measured in a Biotek Synergy HT microtiter plate reader. Data from six wells was averaged for each data point and standard errors calculated. Data for germination and hyphal length assays were collected manually from multiple photographic images taken at 10X and/or 20X.

## Acknowledgements

We would like to thank Dr. Zachary Wood for his instructive discussions about the renaturation and stabilization of sDectin-1 and Dr. Bradley Phillips for discussion on the clinical use of and problems with Amphotericin B. Funding was provided as follows: SA and RBM (University of Georgia Research Foundation, Inc. UGARF, NIAID R21 AI144498, NIH National Center for Advancing Translational Sciences #UL1TR002378), ARF (NSF Graduate Research Fellowship), ZL (RSG-14-184-01-DMC from the American Cancer Society), XL (NIAID 1R01AI140719), MM and SEK (UGA President’s Interdisciplinary Seed Grant Program). None of the authors have any financial conflict of interest.

## Supplemental Tables and Figures

**Supplemental Table S1. Liposome compositions**. Comparison of the chemical composition of liposomes discussed in the manuscript.

**Supplemental Fig. S1. The modified mouse sDectin-1 DNA *MmsDectin1lyshis* and protein MmsDectin-1. A.** The codon optimized DNA sequence of *MmsDECTIN1lyshis* was cloned into in pET-45B. NCBI BankIT submission #2173810. Length: 577 bp, Vector pET-45b sequence highlighted in red with start codon underlined, Cloning sites in green, Codons for Gly Ser (G,S) flexible linker residues in yellow, reactive lys (K) residues in purple, Mouse sDecetin-1 in light blue, terminal Ala codon in yellow to put stop codons in frame, stop codons in bold. **B.** The modified mouse sDectin-1 protein being synthesized. N terminus and His tag from pET-45B vector in red, Gly Ser (GS) flexible linker residues in yellow, reactive lys (K) residues in purple, Mouse sDecetin-1 in light blue. Final Ala residue/codon to put stop codons and PacI site in frame. 199 amino acids, MW 22,389.66 g/mole. Theoretical pI 7.74.

**Supplemental Fig. S2. SDS PAGE analysis of sDectin-1 in cell extracts and after affinity purification.** sDectin-1 protein was produced in the BL21 strain of *E. coli* grown in Luria Broth over night from the pET-45B plasmid without IPTG induction. The protein was solubilized in GuHCl buffers, purified by Ni-NTA resin and examined by SDS PAGE after GuHCl was removed by dialysis. Extraction of protein into buffers that also contained reducing agent beta mercaptoethanol and Triton-X100 detergent greatly increased recovery from insoluble inclusion bodies (center lanes) relative to buffers without them (right lanes). Protein was examined on an 12% acrylamide gel stained with Coomassie Blue. The approximate molecular weight of modified sDectin-1 22 kDa is indicated. Extraction of these cells with urea buffers even at 60oC yielded very little protein (not shown).

**Supplemental Fig. S3. sDectin-coated liposomes, DEC-AmB-LLs, bound strongly to *Candida albicans* and *Cryptococcus neoformans* cells.** A., C. and E. are bright field images of *C. albicans* strain Sc5314 and *C. neoformans* strain H99 labeled with DEC-AmB-LLs diluted 1:100 in LDB. B., D. and F. are the combined bright field and red fluorescence images showing that rhodamine red fluorescent DEC-AmB-LLs bound strongly to these cells. Plain uncoated AmB-LLs and BSA-AmB-LLs did not bind detectably to these cells (not shown). A & B were photographed at 63X under oil immersion, C through F at 20X on an inverted fluorescent microscope.

**Supplemental Fig. S4. sDectin-1 coated DEC-AmB-LLs and BSA coated BSA-AmB-LLs were less toxic to HEK293 cells than uncoated AmB-LLs.** Human Embryonic Kidney HEK293 cells grown to 30 to 40 percent cell density in RPMI lacking red indicator dye in 96 well microtiter plates. Cells were treated for 2 hours with the AmB loaded liposomes indicated or a deoxycholate micelle suspension of AmB (DOC), washed twice and then incubated for additional 16 hrs. All treatments delivered a final concentration of 30 or 15 uM of AmB into the media. The 0 uM control wells received an amount of liposome dilution buffer equivalent to the 30 uM treatment. CellTiter-Blue esterase assays estimated cell viability and survival. Background fluorescence from wells with CellTiter-Blue reagent in the media, but lacking cells and liposomes was subtracted. Standard errors are indicated.

